# Mutation Vulnerability Characterizes Human Cancer Genes

**DOI:** 10.1101/360065

**Authors:** Yong Fuga Li, Fuxiao Xin

## Abstract

Recent studies by Tomasetti et al. revealed that the risk disparity among different types of cancer is mainly determined by inherent patterns in DNA replication errors rather than environmental factors. In this study we reveal that inherent patterns of DNA mutations plays a similar role in cancer at the molecular level. Cancer results from stochastic DNA mutations, yet non-random patterns of cancer mutations emerge when we look across hundreds of cancer genomes. Over 500 cancer genes have been identified to date as the hot spot genes of cancer mutations. It is generally believed that these gene are mutated more frequently because they reside in functionally important pathways and are hence selected during the somatic evolution process of tumor progression. This theory however does not explain why many genes in the same pathways of cancer genes are not mutated in cancer. In this study, we challenge this view by showing that the inherent patterns of spontaneous mutations of human genes not only distinguish cancer causing genes and non-cancer genes but also shapes the mutation profile of cancer genes at the sub-gene level.

## Introduction

Cancer is a group of complex diseases marked by abnormal proliferation of cells as well as somatic mutations of genes. A person has a 40.37% lifetime risk of diagnosis and a 20.84% lifetime risk of dying from cancer (cancer.gov, September 24, 2014). Despite extensive studies, cancer remains a major challenge in medicine. The molecular mechanisms of cancer initialization and progression are not yet fully understood, and for most cancer types, no efficient therapy is available [1].

Cancer causing genes, or cancer genes for short, refers to the class of genes that exhibit causal mutations, either somatic or germline, in cancers [2]. Continued efforts have been devoted to the discovery and study of these cancer genes, with around 15-30 novel ones discovered each year (see supplementary material and **Fig. 1**). Around 550 cancer genes have been identified as of 2016 [2]. These cancer genes, comprising 2.6% of human proteome, span a very wide range of cellular functions and molecular pathways [3], including cell proliferation, growth suppression, DNA repair, apoptosis, cellular senescence, angiogenesis, metastasis, immune response, and energy metabolism. Cancer genes are recognized as the key for understanding the biology of cancer, while cancer gene mutations in individual tumor hold the key for cancer precision medicine.

**Figure 1:**
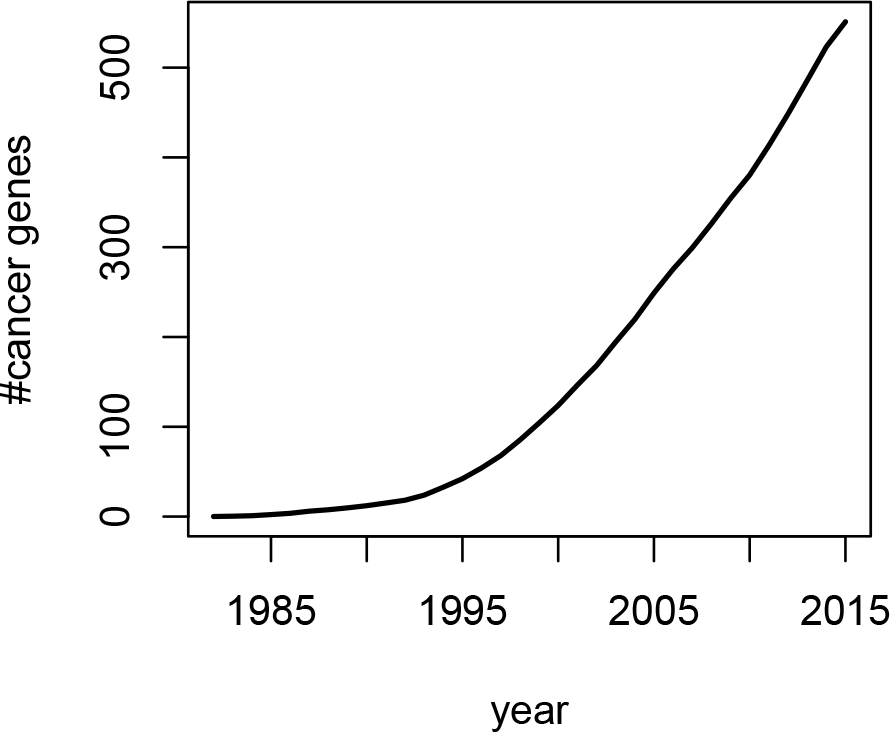
Trend of cancer gene discovery, estimated according to the number of PubMed articles reporting novel tumor suppressor genes or oncogenes.

**Figure 2:**
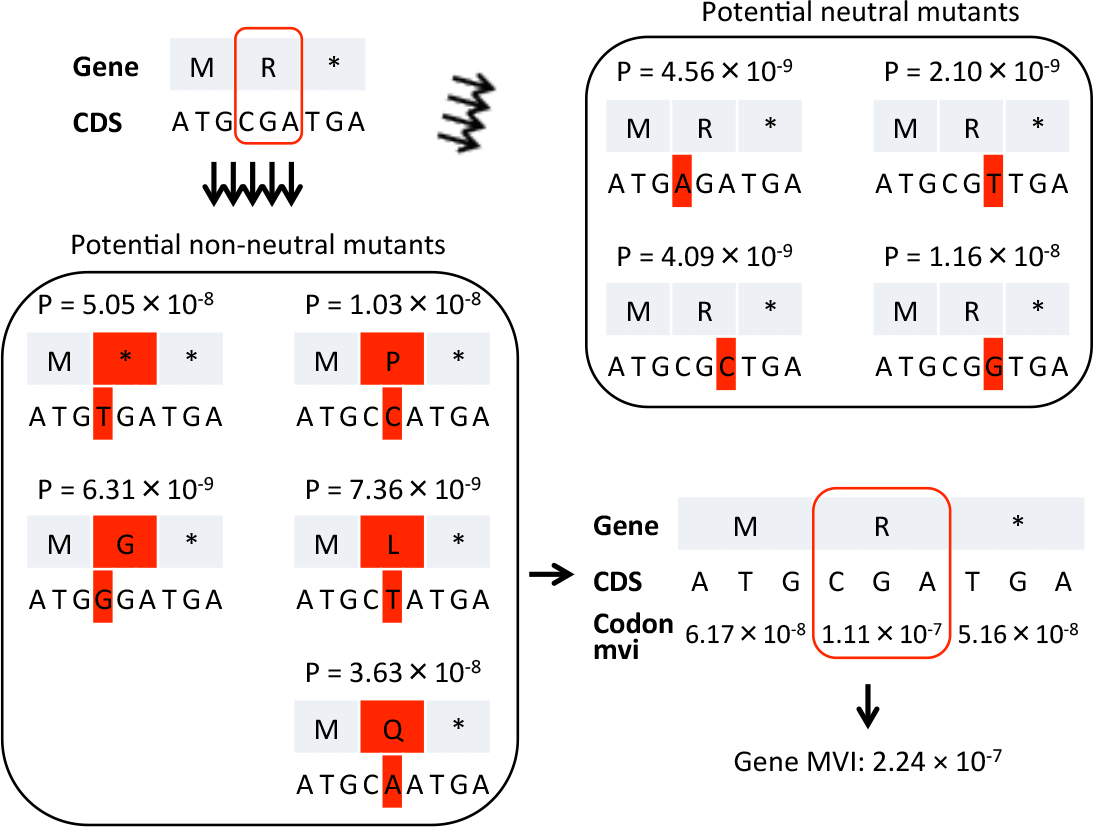
The mutation vulnerability index calculations for a fake gene of three codons. For each codon, there are total nine possible single nucleotide mutations with different probabilities per one round of DNA replication. Di-nucleotide and tri-nucleotide mutations are of low probability and ignored. The nine single nucleotide mutations for the 2nd codon are shown in the figure. Among the nine mutants, five are non-synonymous. Codon mvi values are calculated based on the total probability of these non-synonymous mutants, and gene MVI is the sum all codon mvi values of the gene. Red shades highlight the amino acid and nucleotide mutations.

Despite our successes in discovering cancer genes, the fundamental distinctions between cancer genes and non-cancer genes remain undefined. Studies of the cancer genes repeatedly identify a handful of key biological pathways [3], yet many genes in the same pathways lacks recurrent mutations despite being functionally related to cancer [1, 3, 4]. Relatedly, many cancer genes mutate only in specific type of cancers, or shows drastically different mutation frequency in different types of cancers. These hint that protein function is not the only determinant of cancer genes. Recent studies by Tomasetti et al. revealed that the prevalence difference among different types of cancer is mainly determined by inherent patterns in DNA replication errors rather than environmental factors [5, 6]. In this study we reveal that inherent patterns of DNA mutations plays a similar role in cancer at the molecular level on cancer genes.

Cancer has been recognized as a process of cellular evolution inside human body [7, 8]. In light of the molecular evolution theory, we propose gene’s mutation vulnerability as an independent factor, which together with gene’s specific functions, distinguish cancer genes from other genes. Specifically, different genes are of different spontaneous nucleotide mutation rates, different vulnerability to amino acid changes upon nucleotide mutations, and different vulnerability to functional changes upon amino acid mutations. The vulnerability of a cancer related gene to function impairment sets the stage for somatic selection pressure to play its role.

Based on this theory, we hypothesize that cancer genes are more vulnerable to mutations compared to other genes. To test the hypothesis, we quantify the mutation vulnerability (MVI) of a gene based on the coding DNA sequence (CDS), the DNA mutation rate and spectrum, and the genetic table. We discovered that gene MVI differs significantly between cancer genes and non-cancer genes, and also among different types of cancer genes. Further, we show that MVI of a cancer gene is predictive of its overall mutation frequency in cancers. Finally within individual cancer genes, we show that codon level mutation vulnerability is predictive of the observed codon mutation frequency.

## Materials and Methods

### Mutation vulnerability index

We use a basic probabilistic model to describe the mutation of genes during somatic evolution. Let *CDS* = *N*_1_*N*_2_…*N_j_…N_L_* be a coding sequence of length *L*. We model the probability of nucleotide *N_j_* mutating to 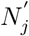 after one round of DNA replication as *P* (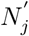*|CDS*) = 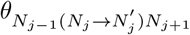, where 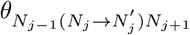 is the neighbor-dependent single nucleotide mutation rate [9, 10]. Notice that, 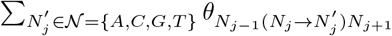 = 1 for any given *N_j_*. In this study, we only model single nucleotide mutations and ignore in-dels and more complex mutation types. Further we assume the nucleotides mutate independently conditioned on the current CDS, i.e. 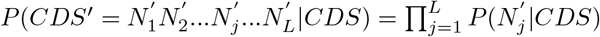. As a result, the probability of two point mutations happening on a single codon or within a single gene is negligible.

We then define the mutation vulnerability index *mvi_x_* for a codon *x* of trinucleotide *N_j_N_j_*_+1_*N_j_*_+2_ as the expected number of nonsynonymous substitution on this codon after one round of DNA replication:

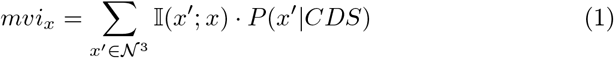

where 𝕀(*x′*; *x*) is the identity function that takes 1 if and only if two codons *x* and *x′* are nonsynonymous, while *P* (*x′|S*) is calculated as *P*(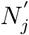|*S*) *· P* (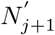|*S*) *· P* (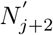|*S*). Notice that 𝕀(*x′*; *x*) measures the functional consequence of mutations, while *P* (*x′|S*) measures the mutational biases. In this study, we use a simple identity function 𝕀(*x′*; *x*) that treats all nonsynonymous mutations as equally non-neutral regardless of the amino acid types and locations in the protein, and similarly treats all synonymous mutations as neutral. *MV I* for a gene is further defined as the expected number of nonsynonymous substitutions on the CDS after one round of DNA replication, and it is calculated as the summation of codon mvi:

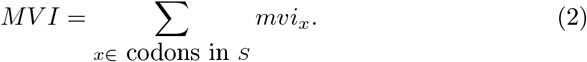

### Predictive models and codon importance

Three types of machine learning methods, including logistic regression, random forest, and neural network, are evaluated with 10 fold cross-validation to determine their performances in predicting cancer genes. Random forest and neural network are evaluated here in order to detect the potential contributions of codon interactions to cancer gene status. Specifically, the data were randomly divided into 10 folds, and 10 separate models were each trained with 90% of the data and tested on the remaining 10%. For training of random forest models, the same number of non-cancer genes were randomly sampled to match the size of the cancer genes in the training set, and 5000 trees were grown. For each machine learning method, five set of models were built using different feature variables: *Len*, only the CDS length is used in building the models; *mvi*, only the precomputed average codon mvi for each gene is used; *Len + mvi*, both CDS length and average codon mvi are used; *codon*, the 64 codon frequencies for a gene are used; *Len* + *codon*, both CDS length and the 64 codon frequencies (normalized by length) are used.

Two measures were used to quantify the importance of each type of codon (feature) in predicting cancer genes: 1) the coefficients of the codon frequency variables in the generalized linear model with logit link function; 2) the *mean decrease of classification accuracy* associated with each codon in the random forest model.

### Mutation rate and spectrum

Due to the chemical properties of DNA as well as cell’s DNA repair machinery, nucleotide’s mutation rate and spectrum vary depending on the identity of the nucleotide, the neighboring nucleotides, as well as the local genomic context [10–12]. For cancer genes, both germline and somatic mutations play important roles. The somatic mutation spectrum can be different from the germline mutations depending the carcinogen that drives the mutations [13, 14], while the overall mutation rate in detectable cancer tissues are generally higher than non-cancer cells [15]. However, the somatic mutation rate and spectrum vary depending on the carcinogen, cancer type, and cancer stage. It is hard to obtain a universal somatic mutation spectrum to model all cancer genes together. Here we use the germline mutation rate and spectrum to model all cancer gene mutations. We believe it is a good model for the germline cancer gene mutations, as well as the somatic cancer gene mutations at the dormant or initiation stage of cancer. The germline mutation rate has been estimated based on mutations of pseudogenes to be around 2.5 *×* 10^*−*8^ [16]. The neighbor-dependent mutation rates are then obtained by adjusting the neighbor-dependent mutation spectrum [10] with this mutation rate. The full mutation rate table is available in Supplementary table 1.

### Gene Sequence and Mutation Data

The CDS sequences and cancer gene mutations are obtained from the Catalogue Of Somatic Mutations In Cancer (COSMIC: http://cancer.sanger.ac.uk/cosmic)[2]. For each gene, only one representative CDS, generally the longest CDS for the gene, is selected in COSMIC to avoid redundancy. This also helps to avoid over-fitting during machine learning. Such strategy is not without limitations. For example, some cancer genes have multiple splicing isoforms of distinct functions in cancer, such as the p16^INK4a^ and p14^ARF^ proteins for CDKN2A gene [17]. Using all 28412 CDS sequences in COSMIC or restricting to Entrez gene (19123 in total) gives nearly identical results. Across this study, only mutations fall in the coding sequences are considered.

## Results

### Human cancer genes show higher mutation vulnerability

There are two prerequisites for a gene to be cancer genes. First, the gene is mutated in cancer cells; Second, specific mutations of the gene renders the host cells with phenotypic changes that allow cancer initiation and progression. To study cancer genes with an emphasis on the mutational aspects in addition to gene functions, we define the mutation vulnerability index (MVI) of a gene as the expected number of nonsynonymous mutations on the gene upon one cycle of DNA replication. We calculated the MVIs of all protein coding genes based on the coding sequences (CDS) and the neighbor-dependent nucleotide mutation rates estimated on human pseudogenes [10]. We then systematically studied the association between gene’s mutation vulnerability and its cancer gene status. We observed a global increase in mutation vulnerability of known cancer genes compare to other genes (**Fig. 3A**). The average gene level MVI of cancer genes is 48.8% higher compare to that of non-cancer genes (p-value 6.9 *×* 10^*−*32^, based on two sample T-test on the log transformed MVIs). We dissected the cancer genes based on whether somatic mutation or germline mutations are observed for them (**Fig. 3B-D**). Cancer genes with only somatic or only germline mutations do not significantly differ in their MVIs, although they have 45% higher MVI than non-cancer genes. On the other hand, cancer genes with both somatic and germline mutations have higher MVIs compare to those with only somatic mutations (p-value 0.03) or only germline mutations (p-value 0.16), or neither (p-value 1.0 *×* 10^*−*8^). We further dissected the cancer genes based the molecular genetic of the mutations. Cancer genes with only dominant or only recessive mutations have 34% or 92% higher MVIs compared to non-cancer genes (p-values 4.8 *×* 10^*−*^21 and 1.3 *×* 10^*−*^13 respectively), while recessive cancer genes have 58% higher MVIs compared to dominant cancer genes (p-value 1.9*×*10^*−*5^). Cancer genes with both dominant and recessive mutations have 206% higher MVI compared to the non-cancer genes (p-value 0.0004).

**Figure 3:**
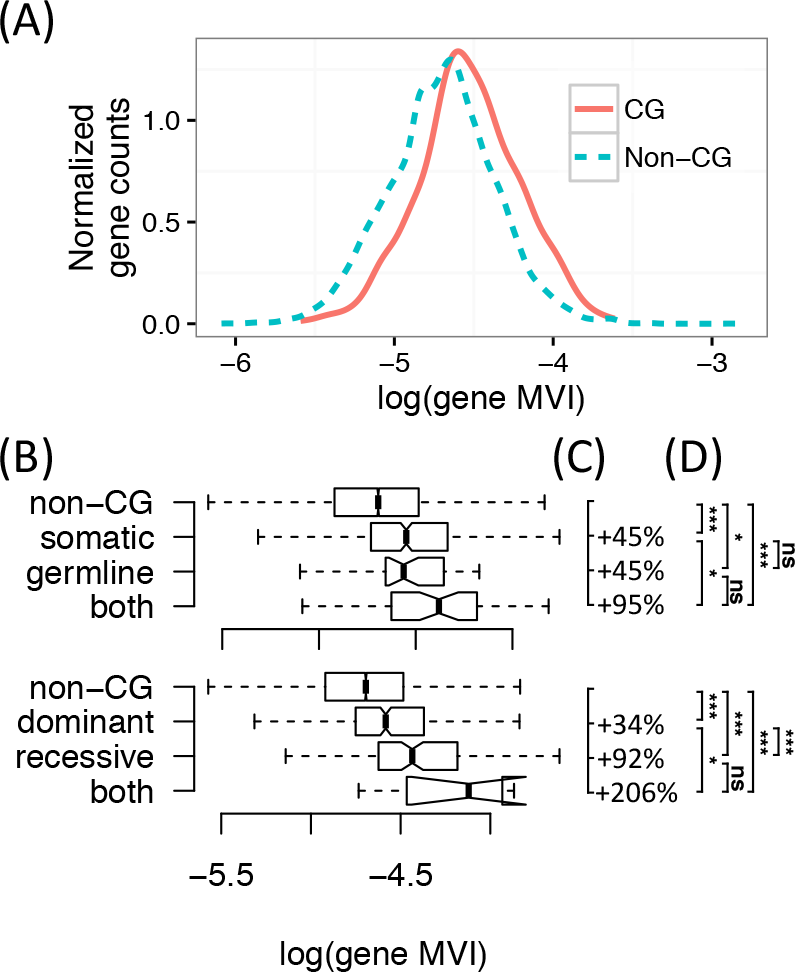
Gene’s mutation vulnerability is associated with its cancer gene status and type. A. Distributions of the gene level MVI of cancer genes (CG) versus non-cancer genes (non-CG). Log base 10. B. Bar plots comparing the MVI of different types of cancer genes against non-cancer genes. On the top: dominant, cancer genes with dominant mutations, mostly oncogenes; recessive, cancer genes with recessive mutations, mostly tumor suppressor genes; both, cancer genes with both dominant and recessive mutations. On the bottom: somatic, genes with only somatic mutations; germline, cancer genes with only germline mutations; both: cancer genes with both somatic and germline mutations. Log base 10. C. The increase in mean MVI of different types of cancer genes relative to that of non-cancer genes. D. Significance levels of pairwise differences of mean MVI among different types of cancer genes and non-cancer genes. Pairwise T-tests were used to compare Log transformed gene MVI. P-values are adjusted for multi-testing using Benjamini–Hochberg’s method. *, p 0.05; **, p 0.01; *** p 0.001; ns: not significant.

### Both protein length and codon mutation vulnerability are predictive of cancer gene

Coding sequence (CDS) length is a major contributor to mutation vulnerability. Let *mvi*_0_ and *mvi*_1_ be the lower and upper bound of the *mvi* for a single codon. The MVI of a gene is then bounded by *mvi*_0_ *· L* and *mvi*_1_ *· L*, where *L* is the length of the protein. The neighbor-dependent nucleotide mutation rate is in the range of 1.2 *×* 10^*−*9^ to 6.1 *×* 10^*−*8^ (Supp. Table 1), and based on this mutation rate, the codon *mvi* is estimated to be in the range of 2.4 *×* 10^*−*8^ and 1.4 *×* 10^*−*7^. For human genes, the length of coding sequences explains 99.81% of the variances of gene *MVI*, while the average codon *mvi* only explains 0.29%. We therefore decompose *MVI* into gene length and average codon *mvi* to study their respective contributions to the *MVI*-cancer gene stustus association.

Gene CDS length alone is associated with cancer genes. Cancer genes are 644 amino acids on average, 46% longer than non-cancer gene which are 441 amino acids on average (p-value 4.4 *×* 10^*−*30^, Wilcoxon rank sum test). Is this cancer gene-CDS length association biological? Here is a counter argument. COSMIC generally selects the longest RefSeq cDNA sequence as the representative sequence (personal communications), as a result, genes with more known isoforms will tend to have slightly longer CDS due to the selection bias. This could lead to artificial associations between CDS length and cancer genes, because cancer genes are generally more extensively studied experimentally compared to non-cancer genes, and may as a result have more isoforms identified, and hence longer longest CDSs. To clarify this, we systematically analyzed gene isoforms based on human RefSeq sequences. Indeed, a cancer gene has 7.01 isoforms on average, compared to 5.03 for non-cancer genes (p-value 4.1 *×* 10^*−*21^, Wilcoxon rank sum test). To eliminate the biases associated with isoform counts, we examined the median rather than max length of each gene’s isoforms, and found that cancer genes are still 45% longer than non-cancer genes (591AA versus 407AA, p-value 2.2 *×* 10^*−*28^). Even the shortest isoforms of cancer genes are 35% longer than the shortest isoform of non-cancer genes (454 AA versus 337 AA, p-value 1.0 *×* 10^*−*18^), despite that we expect the lengths of the shortest isoforms to be negatively biased by intense research activity on cancer genes. Statistical evaluation further confirms the conditional dependence of cancer gene status with CDS length after controlling for the number of isoforms (Jonckheere-Terpstra Test on ordinal variables, *z* = 6.21, p-value = 5.2 *×* 10^*−*10^). These observations hence provide strong evidence that the CDS length-cancer gene status association is biologically real.

Although CDS length is the main contributor of gene’s mutation vulnerability, gene’s average codon *mvi*, calculated as *MV I/L*, remain significantly associated with cancer genes (p-value 0.0005, Wilcoxon rank sum test). The association remain significant after controlling for CDS length (Jonckheere-Terpstra Test, *z* = 4.20, p-value = 2.7 *×* 10^*−*5^). With these, we suggest that both CDS length and codon usage are biologically associated with cancer genes, with CDS length being the main contributor to the association.

### The predictive power of codon usage on cancer genes

The mutation vulnerability index calculated based on Equation 2 has two limitations. First, it relies on the germline nucleotide mutation spectrum, which was derived based on pseudogenes [10]. Second, it utilizes a naive model for amino acid mutation effect such that all nonsynonymous mutations lead to protein function impairment. One way to overcome these limitations is to use machine learning and directly model the relationship between the codon usage and cancer gene status. We refer to the learned probability from such data-driven model as the *learned MVI* as compared to the *precomputed MVI* from Equation2.

Using the normalized codon frequencies together with protein length as predictor variables (*Len* + *codon*) in logistic regression, we achieved an AUC (area under the Receiver Operator Characteristic curve) of 0.731 (95% confidence interval 0.710-0.751, estimated from 30 times 10-fold cross validations). If only the 64 normalized codon frequencies (model *codon*) are used, the AUC is 0.727 (95% confidence interval 0.707-0.747, **Fig. 4** and Supp. **Fig. S2**). For comparison, precomputed *MVI*, protein length (*Len*) and average codon *mvi* alone each achieved AUC 0.646, 0.641 and 0.543 respectively, while AUC 0.661 is achieved when protein length and average codon mvi are combined through logistic regression (*Len* + *mvi*). The superior predictive power of learned MVI from models *Len* + *codon* and *codon* compared to the precomputed *MVI* support the limitations of the germline mutation spectrum and naive mutation effect model. Similar predictive performances are observed when random forest or artificial neural network are used instead of the logistic regression (data not shown), suggesting that nonlinear combinations of the codon frequencies do not contribute to cancer gene status. Notice that the precomputed MVI considers the impact of two neighboring nucleotide of a codon on the mutation rate of the 1st and 3rd nucleotides in a codon, while the machine learning models do not captures such information.

**Figure 4:**
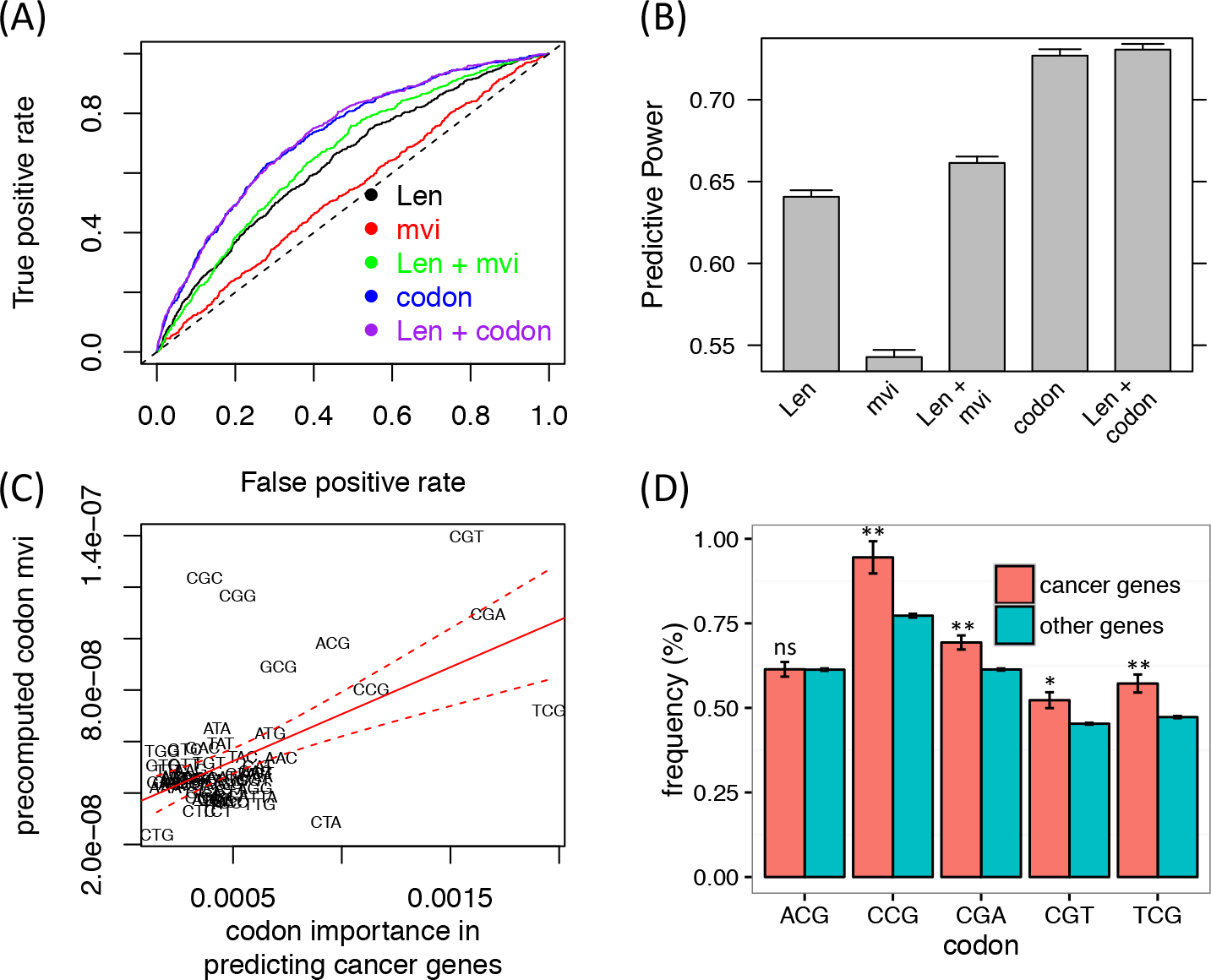
The power of gene’s CDS length (*Len*), average codon MVI (*mvi*, i.e. *MV I/Len*), and codon usage (*codon*) in differentiating cancer genes from non-cancer genes. A) Receive Operator Characteristic curve for representative 10-fold cross validations for models *Len*, *mvi*, *Len* + *mvi*, *codon*, and *Len* + *Codons*. B) The performance comparison among the 5 models. The error bars are estimated based on 30 iterations of 10-fold cross validations. C) Importance of codons in predicting cancer genes is significantly associated with the mutation vulnerability of the codons. Only non-stop codons are shown. Pearson correlation 0.55, p-value 2.0 *×* 10^*−*6^. (D) The frequencies of top 5 most important codons in cancer genes versus non-cancer genes. * p-value < 0.01, ** p-value < 0.001.

### Learned codon importance recapitulates precomputed codon mutation vulnerability

Based on the machine learning models, we analyzed the importance of each of the 64 codons in predicting cancer genes. Interestingly, we observed that codons of higher importance in predicting cancer genes also have higher precomputed codon *mvi*, regardless of the machine learning models (generalized linear model or random forest). In another word, the relative importance of the codons recapitulates codon’s mvi. If mutation vulnerabilities of codons do not impact genes’ cancer gene status, we do not expect to observe this relationship.

For generalized linear model, we measure the importance of each codon by its coefficient in the model fitted either 1) with all codons together or 2) with each codon separately. For the former, we observed a positive Pearson correlation coeffcient 0.26 (p-value 0.021) for 61 non-stop codons, and correlation coefficient 0.12 (p-value 0.18) for all 64 codons (Notice that the stop codons are not very informative since each gene has only 1 stop codon.) For the latter, we observed a positive correlation 0.55 (p-value 2.0 *×* 10^*−*6^) for the non-stop codons and correlation coefficient 0.42 (p-value 0.00032) for all 64 codons (**Fig. 4C**). Similar results are obtained for random forest with *mean decrease of classification accuracy* as codon importance measure.

The 5 most importance codons are TCG, CGA, CGT, CCG, and ACG. Notice they all contain CpG di-nucleotide, which are known to be associated with high mutation rates due to methylation. Among these five codons, four have significantly higher codon usage in cancer genes compared to non-cancer genes (**Fig. 4D**), with 21%, 13%, 15%, and 22% higher frequencies in cancer genes for codons TCG, CGA, CGT, CCG respectively.

It is worth noting that across human CDSs, there is a global negative association between the codon’s mutation vulnerability and the codon usage (Pearson correlation *R* = −0.37, p-value = 0.0026, or *R* = -0.43, p-value = 0.00048 after removing the stop codons, **Supp. Fig. 1**). This is consistent with the mutational bias theory of codon usage biases, which states that the codons that are easily mutated at the nucleotide level to other codons will end up having low frequency.

### The variation of MVI among cancer genes is associated with their mutation frequency

While we observed that gene *MVI* differs between cancer genes and non-cancer genes (**Fig. 3-4**), we also notice large variation of *MVI* among cancer genes themselves (**Fig. 3**). Are the *MVI* differences between two cancer genes of any biological significance?

We believe *MVI* together with gene’s function determine the probability that a gene is observed mutated in cancer. There are two predictions from this: first, MVI will impact a gene’s cancer gene status; second, MVI will impact the mutation frequency of cancer genes. We have evaluated and confirmed the first prediction. To validate the second prediction, we estimate each cancer gene’s mutation frequency in cancer samples based on the mutation data in COSMIC. Notice that not all samples in COSMIC were subjected to full genome sequencing, hence the estimation can be conservative and inaccurate. Despite this, we observed significant association between cancer genes’ MVIs and their mutation frequencies (**Fig. 5**). There are significant positive associations between precomputed *MVIs* and mutation frequencies for tumor suppressor genes (Spearman rank correlations 0.687, p-value 1.4 *×* 10^*−*19^) as well as for oncogenes (Spearman rank correlations 0.580, p-values 8.7 *×* 10^*−*37^). Similarly, machine learned MVI is also positively correlated with mutation frequencies for tumor suppressor genes (Spearman rank correlations 0.444, p-value 1.1 *×* 10^*−*7^) as well as oncogenes (Spearman rank correlations 0.105, p-values 0.04). We emphasize that the learned MVI is trained for differentiating cancer gene against non-cancer gene without any knowledge of cancer gene’s mutation frequency, yet it is significantly associated with the mutation frequency of cancer gene in tumor samples. These further validate the association between mutation vulnerability and cancer genes.

**Figure 5:**
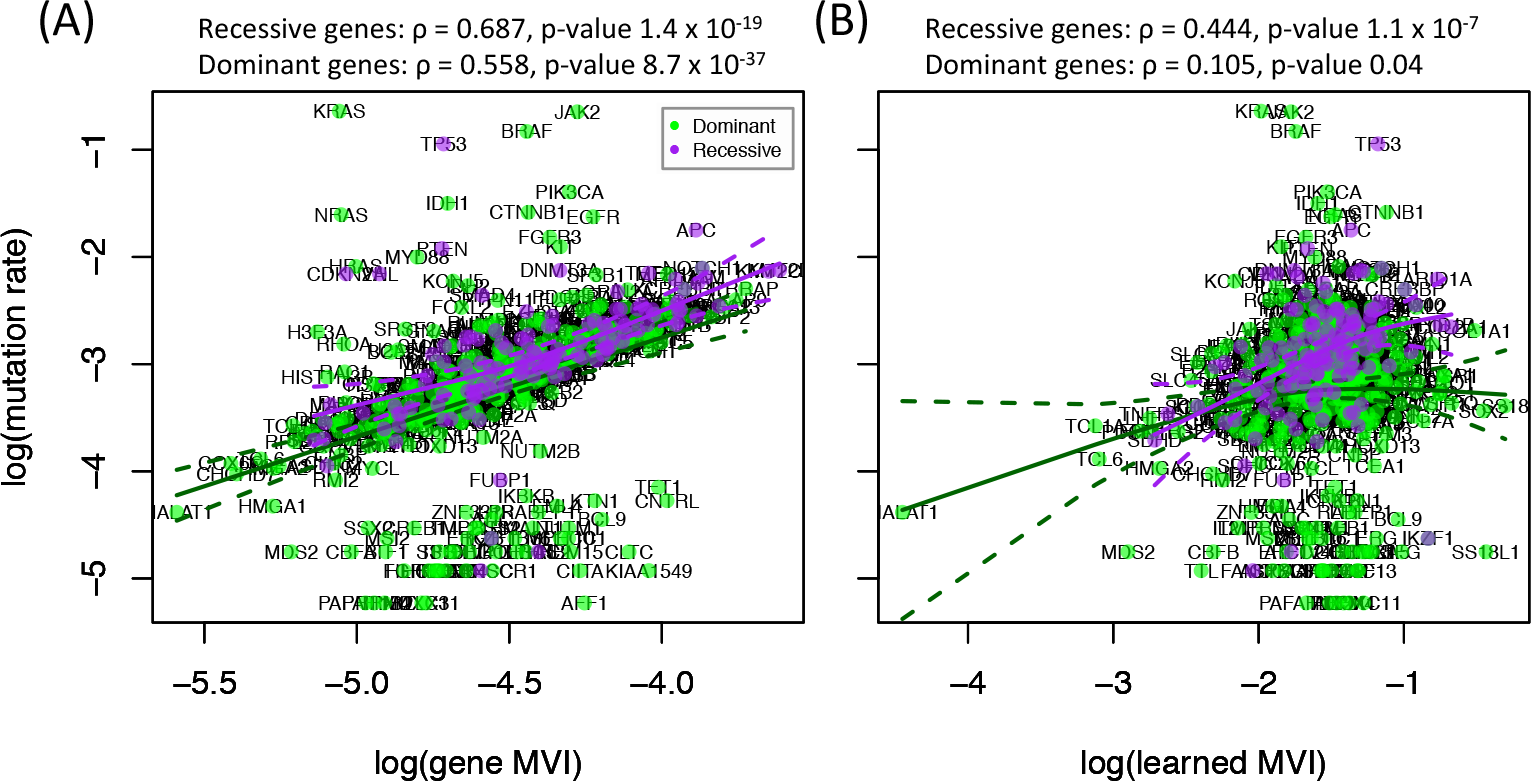
A cancer gene’s MVI is positively associated with its mutation frequency in cancer samples. Scatter plot of the mutation frequencies of cancer genes against precomputed MVI (A) or learned MVI (B). In (B), the learned MVI is trained using the codon usage as predictors and gene’s cancer gene status as target, hence the mutation frequency information is completely independent from the machine learning procedure for cancer gene prediction. Gene’s mutation frequencies in cancer samples are calculated based on COSMIC gene mutation database. Log base 10 is used.

### Intra-gene codon mutation frequency and codon mutation vulnerability are associated

Cancer is a process of somatic evolution. If somatic selection pressure is uniformly positive for all amino acid mutations in cancer proteins, then the spontaneous mutation rate of an amino acid decides its mutation frequency in cancer, and the spontaneous mutation rate of a cancer protein decides the cancer gene’s mutation frequency in cancer. We have studied the gene level mutation frequency in the previous section, and here we study the codon level mutation rate.

For each of the 541 cancer genes with mutation data, we determine if the observed mutation frequency of the codons (along the CDS) are correlated with precomputed codon mutation vulnerability (*mvi*). Positive linear correlations are observed for 89% (482) of the genes, among which 71% (342) are significant at p-value cutoff 0.01, and 52% (253) remain significant after Benjamini-Hochberg correction. By contrast, among the 59 genes with negative correlations between *mvi* and observed codon mutation frequency, none are significant at p-value cutoff 0.01 (**Fig. 6A**). The codon mvi-codon mutation frequency correlation is on average stronger for tumor suppressor genes compared to oncogenes (p-value 1.0*×*10^*−*8^, Mann-Whitney test on the ranks of p-values for codon mutation frequency-mvi correlation). 97% of the tumor suppressor genes has positive codon mvi-mutation frequency correlation, 81% among which are significant at p-value level 0.01, while only 87% of oncogenes has positive mvi-mutation frequency correlation, 67% among which being significant. We visualized RB1 and PHF6 as examples to further understand the codon mvi-codon mutation frequency relationship. By plotting the predicted codon mvi and observed codon mutation frequency spectra as mirror images (**Fig. 6B**), we find that mutation vulnerability explains a significant portion of the mutation hot spots. Majority of mutation hot spots have high mutation vulnerability, although many codons with high mvi are not highly mutated, suggesting that high mutation vulnerability is a precondition for high mutation frequency. These corroborate with previous findings and support the important roles of mutation vulnerability on cancer gene mutation.

**Figure 6:**
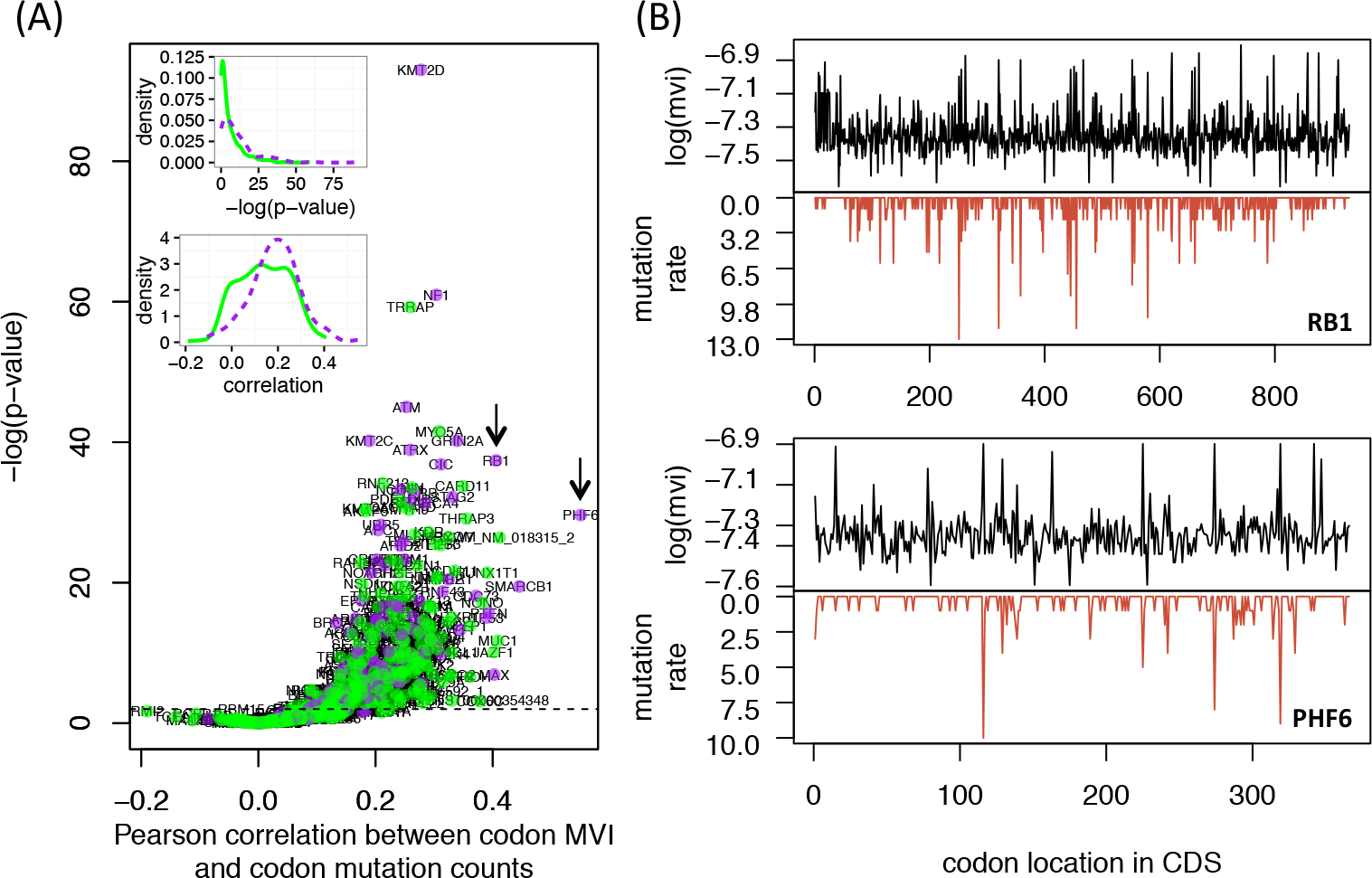
Associations between intra-gene codon mutation frequencies and codon’s mutation vulnerability. (A) Volcano plot for the codon mutation frequency - codon mvi correlation. X-axis is the Pearson correlation coefficient per cancer gene. Y-axis is the p-value associated with the correlation coefficient. Dashed horizontal line marks p-value cutoff 0.01. Inlets: the density plot for correlation coeff-cients (top) and p-values (bottom) grouped by cancer gene type (purple, tumor suppressor genes; green, oncogenes). Arrows point to RB1 and PHF6 genes, for which detailed mutation spectrum and predicted mvi spectrum are shown as mirror images in (B) and (C). The mutation frequency spectra are flipped vertically. Codon mvi is calculated by aggregating the neighbor dependent nucleotide mutation frequencies with the mutation consequence determined by the generic genetic table. Codon mutation frequency is taken as the raw counts of mutations occurring at a given codon. Log base 10 is used.

## Discussion

Cancer is a complex genetic [18, 19] and aging disease [20]. Recently big data approach to cancer genomics study is driving an increasingly complete understanding of the genes and pathways underlying cancer development [21–24], while at the same time revealing the stochastic nature of cancer due to somatic evolution [7, 8]. Over 500 cancer genes have been identified with recurrent (driver) mutations in cancer samples, yet no cancer gene is found to mutate in all cancers, and no two cancers share the same set of cancer gene mutations.

The specific functions, e.g. biological processes, pathways and protein interactions, of genes are believed to be the reason why some genes are cancer genes while others are not [25]. In this study, we analyze the mutational aspect of cancer genes. We found that biases in the DNA mutation process, captured by each gene as its mutation vulnerability (MVI), significantly impact the likelihood for a gene to be observed as cancer gene. Compared to non-cancer genes, cancer genes have on average 48.8% higher mutation vulnerability (**Fig. 3**), with contributions from both protein length and codon usage. Importantly, among the cancer genes, the ones with higher MVI also have higher mutation frequency in cancer samples (**Fig. 5**). This suggests that MVI influences a gene’s chance of being a cancer gene in a quantitative manner. Recent studies [5, 6] revealed that the cancer risk disparity among different tissue types is mainly determined by inherent properties of DNA replication errors related to cell types rather than environmental factors. In this study we reveal that inherent patterns of DNA mutations may play a similar role at the molecular level.

At the codon level, we observe that majority of highly mutated codons in cancer genes have high *mvi* (codon mutation vulnerabilities), while many codons with high predicted *mvi* are not highly mutated (**Fig. 6**). High mutation vulnerability and functional relevance are likely two independent prerequisites for a gene to be cancer gene and for a codon of a cancer gene to mutated in cancer. The mutation vulnerability of genes and codons set the stage for positive selection to play its role in cancer formation [26]. A gene with cancer relevant function but low vulnerability may not show statistically significant mutation frequency to be identifiable as a cancer gene. We suggest that cancer gene should not be viewed as a binary concept but rather a continuum. With more cancer genomes sequences, we will have more power and detect more “weak” cancer genes.

A machine learning approach for predicting cancer gene will help us to understand the key features differentiating cancer gene from other genes, and it may also guide the discovery of new cancer genes and improve the interpretation of cancer genomes in differentiating driver mutations from non-driver mutations. Existing effort for cancer gene prediction has been focused on utilizing the functional attributes of genes, e.g. molecular functions and signaling pathways, locations and connectivity in the protein-protein interaction network [25, 27]. Some recent studies also suggested the value of gene expression [25, 26, 28]. We reveal here that cancer genes can be predicted at a decent performance by precomputed or learned MVIs, both of which are functions of gene’s codon frequencies (**Fig. 4**). Further improvement of cancer gene prediction will be possible if the protein functions and expression levels are combined with the mutation vulnerabilities of genes.

We note that the mutation vulnerability index is inversely related to concept of mutational robustness in evolutionary biology [29, 30], which describes the property that an organism’s fitness remains unchanged upon mutations. The main feature of MVI is that it captures the spontaneous mutation biases of nucleotides as well as the consequences of the nucleotide mutations. MVI, however, does not accurately model if an amino acid change is functionally neutral to the cell or organism.

There are several limitations in our approach for computing gene’s mutation vulnerability. First, we only model single nucleotide mutations. In-dels and rearrangements, which are common for oncogenes, are not considered. Explicit modeling of these mutation types could give us a better mutation vulnerability index for genes. We also only consider the impact of two immediate neighbors on the mutation rate of a nucleotide. Including more neighboring nucleotides as well as the local genomics environment around a gene, e.g. chromatin structure and CG%, could improve the accuracy of MVI. Finally, for the precomputed MVI (Equation 2) non-sense and missense mutations are treated equally, and all missense mutations are viewed the same in terms of their functional impacts. Ongoing research shows that we can better predict the functional consequences of amino acid changes by incorporating the amino acid conservation patterns and neighboring amino acid sequences [31–33]. Using these advanced models instead of the naive mutation effect model could lead to a better MVI, improved cancer gene predictions, and increased understanding of cancer genes.

## Declarations

### Ethics approval and consent to participate

Not applicable.

### Consent for publication

All authors agreed to the publication of the manuscript.

### Availability of data and material

Relevant data are provided as supplementary material.

## Authors’ contributions

YFL participated in the study design, analysis, and wrote the manuscript. FX carried out analysis and revised the manuscript.

## Acknowledgements

The authors would like to thank Tara Brock and Janelle Marie Estes for discussions.

